# The neuronal basis of insect stereopsis

**DOI:** 10.1101/395939

**Authors:** Ronny Rosner, Joss von Hadeln, Ghaith Tarawneh, Jenny C. A. Read

## Abstract

A puzzle for neuroscience - and robotics - is how insects achieve surprisingly complex behaviours with such tiny brains^1,2^. One example is depth perception via binocular stereopsis in the praying mantis, a predatory insect. Praying mantids use stereopsis, the computation of distances from disparities between the two retinas, to trigger a raptorial strike of their forelegs^3,4^ when prey is within reach. The neuronal basis of this ability is entirely unknown. From behavioural evidence, one view is that the mantis brain must measure retinal disparity locally across a range of distances and eccentricities^4–7^, very like disparity-tuned neurons in vertebrate visual cortex^8^. Sceptics argue that this “retinal disparity hypothesis” implies far too many specialised neurons for such a tiny brain^9^. Here we show the first evidence that individual neurons in the praying mantis brain are indeed tuned to specific disparities and eccentricities, and thus locations in 3D-space. This disparity information is transmitted to the central brain by neurons connecting peripheral visual areas in both hemispheres, as well as by a unilateral neuron type. Like disparity-tuned cortical cells in vertebrates, the responses of these mantis neurons are consistent with linear summation of binocular inputs followed by an output nonlinearity^10^. Additionally, centrifugal neurons project disparity information back from the central brain to early visual areas, possibly for gain modulation or 3D spatial attention. Thus, our study not only proves the retinal disparity hypothesis for insects, it reveals feedback connections hitherto undiscovered in any animal species.

---

In humans, stereopsis is supported by a complex network spanning multiple cortical areas and involving tens of millions of neurons^8,11^. Praying mantids achieve stereopsis with brains orders of magnitude smaller than vertebrates’. Thus, it is natural to assume that insect stereopsis must be computed in a profoundly different and much simpler manner^7^. Insect stereopsis does differ from humans’ in using changes in luminance, rather than luminance directly^12^. However, this does not explain how the mantis brain combines information about the location of luminance changes in the two eyes. In primates, individual retinotopic neurons in the primary visual cortex are tuned to different disparities and thus different locations in 3D-space, but such local computations are often regarded as far too neuronally expensive for an insect brain^7,9^. One theory is that prey is identified in each eye separately and this monocular information is combined in a single late stage in the motor pathway ^6,9^. This would require no disparity-sensitive circuitry in the brain.

To determine whether neurons tuned to binocular disparities exist in the mantis brain, we recorded intracellularly in the *optic lobes*, the major visual processing centres in insects (Fig. 1a,b). Animals viewed a computer screen through coloured filters enabling us to control stimuli to each eye separately and thus presenting images in 3D^3^ (Fig. 1a). Neurons were stained for subsequent identification.

**Figure 1.**
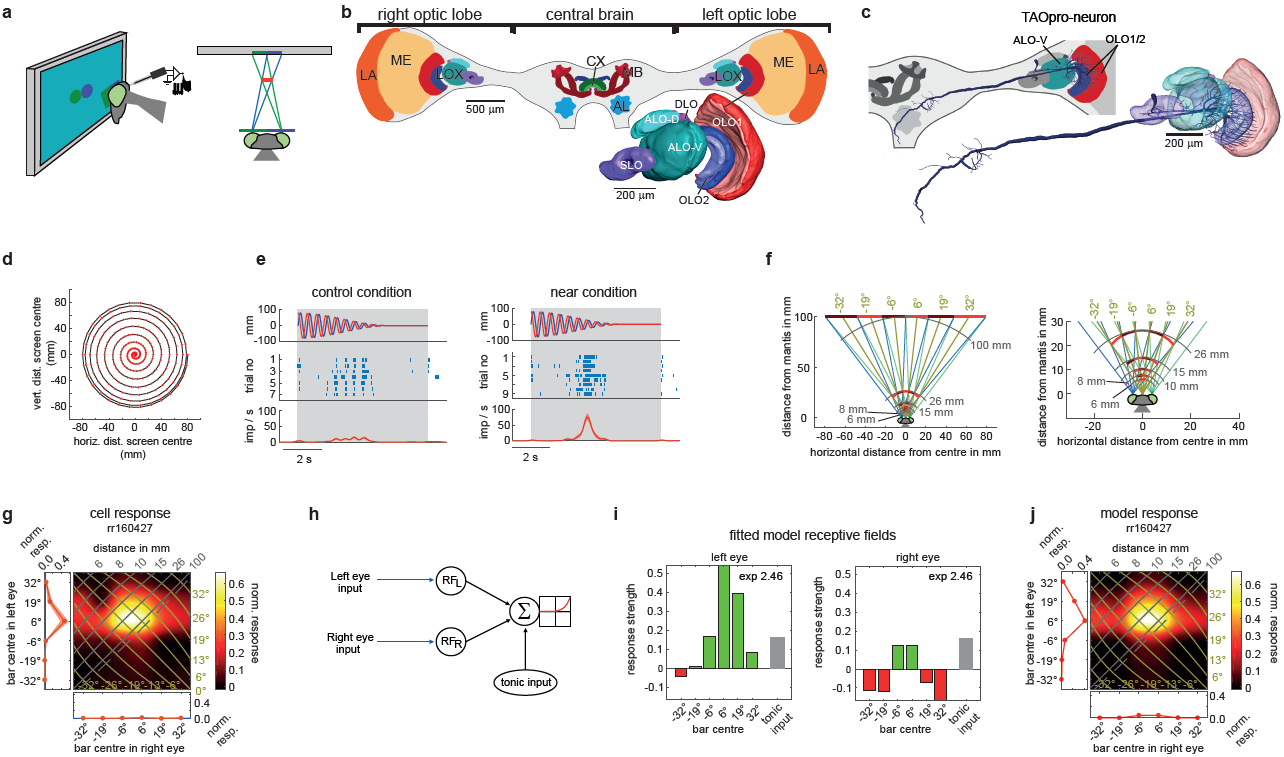
Experimental setup, disparity-sensitive neuron and fitted model. **a**, Praying mantis watches computer screen showing disc stimulus during neuronal recording (side view, top view). Spectral filters ensure each eye sees only one disc; lines of sight in blue (green) for left (right) eye. Virtual disc floats in front of screen (red line). **b**, Mantis brain frontal view with major neuropils. Inset 3D-reconstruction of *lobula complex* (LOX), site of ramifications of all except one neuron type presented in this study. AL, antennal lobe; ALO-D, dorsal unit of the anterior lobe; ALO-V, ventral unit of anterior lobe; CX, central complex; LA, lamina; MB, mushroom bodies; ME, medulla; OLO1/2, outer lobe 1/2; SLO, stalk lobe. **c**, Reconstruction of TAOpro-neuron (tangential <u>pro</u>jection neuron of the <u>a</u>nterior and <u>o</u>uter lobes) showing ramifications in LOX and central brain. Inset: left brain frontal view, colour shows LOX sub-compartments containing ramifications. **d**, Disc trajectory. Red dots: disc centre at 60 Hz refresh rate. **e**, Responses of TAOpro-neuron to disc, visible during grey-shaded region. Upper lanes: vertical (blue) and horizontal (red) distance of disc from screen centre as function of time; negative values are left and lower side of screen. Middle lanes show raster plots, lower lanes spiking rates (average red line, ±1SEM ribbons) after Gaussian smoothing with SD of 150 ms. Right plot: virtual disc at 25 mm (in catch range), left: control (right and left eye stimulus swapped). **f**, Bar stimulus configuration with all 6 different bar positions on computer screen (shown alternating dark/bright red for clarity; right panel zoom) and virtual bars corresponding to different left/right pairs. Azimuthal direction of simulated bars from mantis head midline in brown and distance isolines in grey. **g,** Monocular (marginal, red) and binocular (pseudocolor) responses to bar. Axes show centre of bar in left and right eye. In monocular plots, ribbon shows ±1SEM; blue line shows background activity. Binocular response is interpolated (raw plot in Extended Data Fig. 2). Isolines indicate azimuth (brown) and distance (grey) of virtual bar from mantis as shown in **f**. Dashed line marks screen locations implying objects at “infinity” (parallel lines of sight). **h**, Outline of *Linear-Nonlinear Model* components. Visual inputs are filtered by left-and right-eye receptive fields, then summed linearly along with tonic input, followed by spiking threshold and power-law. **i,** Fitted left and right receptive field functions for TAOpro-neuron; green = excitation, red = inhibition, grey = tonic input (shown in both RF plots, but applied only once), exponent in upper right corner (exp). **j,** Fitted model responses for TAOpro-neuron.

A tangential projection neuron of the optic lobe, “*TAOpro*”, is well suited to detect stereoscopically-defined mantis prey. It ramifies in the outer lobes and the anterior lobe of the lobula complex (LOX), a highly structured visual neuropil in the mantis brain^13^ (Fig. 1c). We recorded TAOpro’s responses to a “spiralling disc” stimulus (Fig. 1d) which mimics mantis prey. During behavioural experiments mantids readily strike at the disc when its disparity indicates it is in catch range, but not in the control condition with reversed disparity^3^. When the same stimulus was presented to the restrained praying mantis during neuronal recording, the TAOpro-neuron responded vigorously for the disparity indicating catch range, and only weakly for the control condition (Fig. 1e; Wilcoxon rank sum test, p=5.2×10^-4^).

To understand the neuronal computation supporting this response, we used our main stimulus comprising vertical bars, 13° wide and flashed briefly at six different, non-overlapping locations independently in each eye (Fig. 1f and Methods). Vertical bars avoided the need to identify receptive field elevation while enabling us to vary horizontal disparity. For studying potential “prey-detector” neurons, we used dark bars on a brighter background, since mantids strike preferentially at dark prey^14^. Each eye saw either a single bar or a blank screen. In this way we simulated virtual objects at a range of 3D locations in front of the animal (Fig. 1f), as well as “control” locations not corresponding to any single location in space (Extended Data Fig. 1).

Tested monocularly, the TAOpro-neuron responded only to bars presented in the left eye (Fig. 1g). However, the single blob-like peak in the binocular response field shows that it receives binocular input. This response resembles disparity-tuned *simple cells* in the mammalian cortex^10^ and means that the neuron responds selectively to a combination of object locations in left and right eyes, and thus to a particular location in 3D-space. The response peaked for left-eye stimulation at 6° and right-eye at 0°, corresponding to an object located ˜3° to the right of midline at a distance of ˜50 mm (cf azimuth/distance isolines Fig. 1f) - an ideal strike target location for mantises^15^. For other locations, the response is much weaker, because input from the right eye provides an inhibitory surround which acts to suppress the neuron’s response to stimuli closer than 15 mm or further away than 100 mm (Fig. 1i, right panel). Similar inhibitory surrounds were found in other neurons (see below).

In vertebrate stereopsis, disparity-tuned simple cells are well described by an influential model^10^ which assumes that each eye’s image is filtered by a linear receptive field, and the result summed and passed through a threshold-plus-power-law non-linearity (Fig. 1h). We fitted this model to our mantis neuron responses. The free parameters were the outputs of the monocular receptive fields in response to a bar at each of the 6 locations in each eye, plus any tonic input and the exponent of the non-linearity (see Methods). This model accounted well for the response of TAOpro (Fig. 1j) and most other neurons we investigated (see below).

The TAOpro-neuron projects to the ventromedial protocerebrum into what corresponds to the *vest* and/or the *posterior slope* in other insects^16,17^. Here descending neurons relaying visual information to thoracic motor centres receive input^18–20^. Thus, TAOpro could directly deliver the signal for the raptorial strike.

Since the TAOpro-neuron receives direct input only in the left optic lobe, this raises the question of how it obtains its information from the contralateral eye. We identified an array of columnar neurons, *COcom*, (Fig. 2a; Extended Data Fig. 3) transmitting visual information from one optic lobe to the other. They possess beaded (i.e. output^21^) ramifications contralateral to the cell’s soma, but smooth (input^21^) endings ipsilaterally (Fig. 2b,c). They also send output fibres to the central brain.

**Figure 2.**
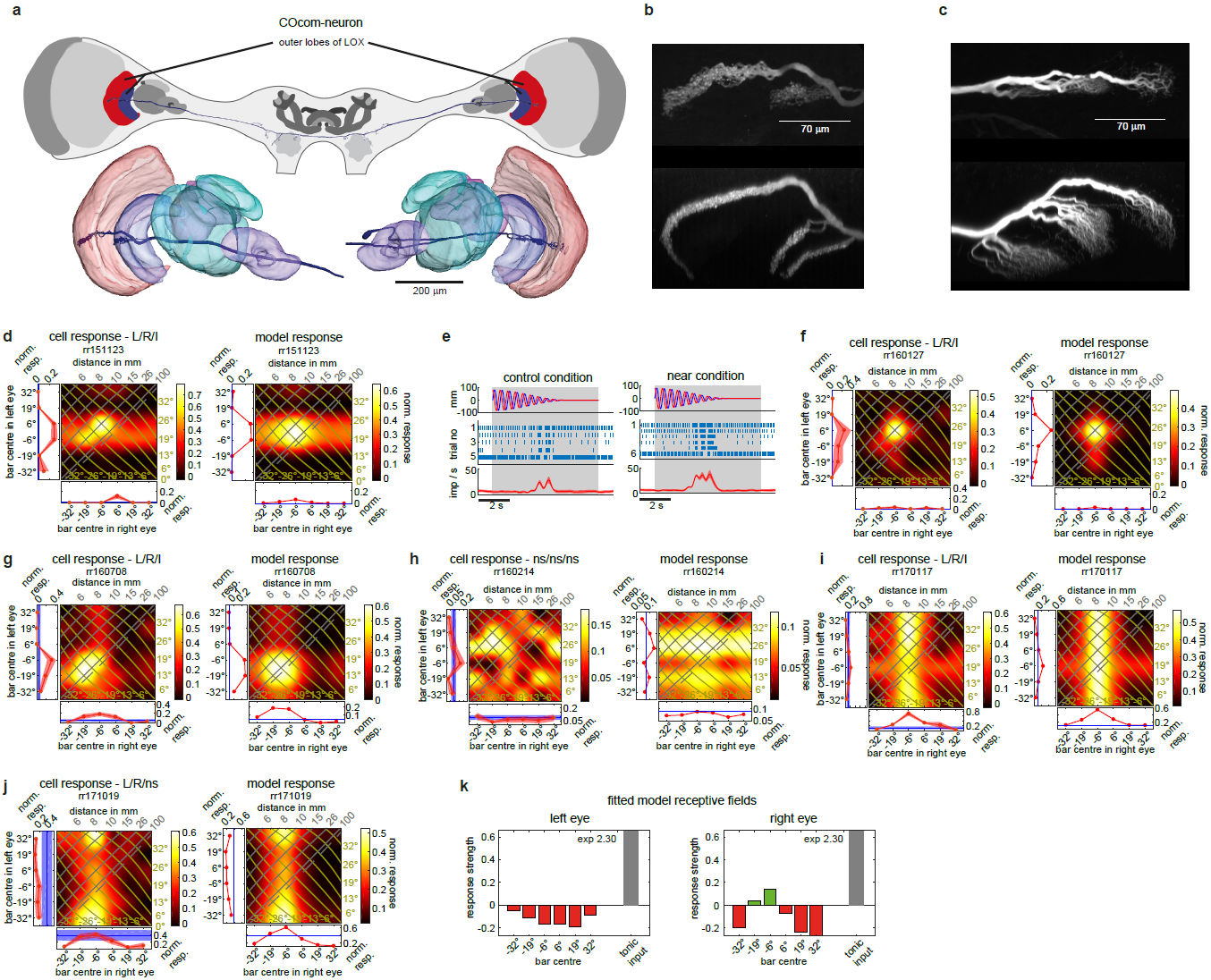
Columnar, commissural neuron, connecting both lobula complexes. **a**, COcom-neuron (<u>c</u>olumnar, <u>com</u>missural neuron of the <u>o</u>uter lobes) reconstruction, anterior view. Neuron ramifies in outer lobes of both LOX and in central brain. **b**,**c**, Maximum projections of confocal horizontal (anterior to posterior orientation; top) and coronal sections (dorsal to ventral orientation; bottom) through outer lobe 1 and 2 in right (**b**) and left (**c**) optic lobe. Terminal neurites are strongly globular/beaded in right (**b**) and smooth in left optic lobe (**c**). **d**, Measured (left) and modelled (right) response fields for neuron in **a, b, c** for flashed dark bars. **e**, response of same neuron to spiralling disc. **f-j** as **d** for 5 further COcom-neurons. Response field headers provide outcome of two-way-ANOVA with “L” (“R”) being significant left (right) eye input and “I” significant interaction term (see Extended Data Table 1), otherwise “ns” meaning not significant. **k**, Fitted receptive fields for neuron in **j**.

**Figure 3.**
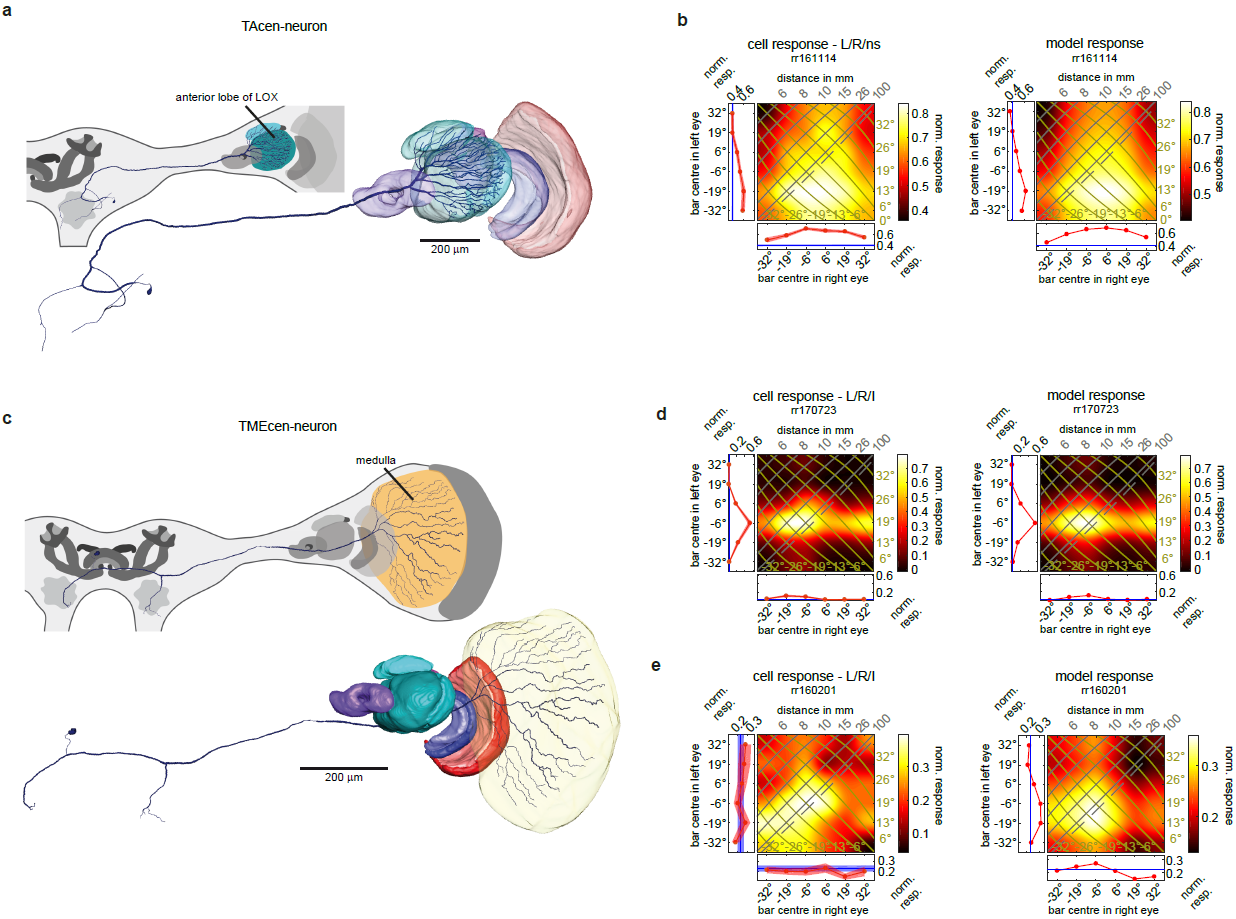
Centrifugal (feedback) neurons. **a,** Anterior view of reconstructed TAcen-neuron (tangential <u>cen</u>trifugal neuron of the <u>a</u>nterior lobe) with ramifications in anterior lobe of LOX and central brain. **b,** Cell (left) and model (right) response fields for a representative TAcen-neuron to flashed, bright bars (responses of 5 further TAcen-neurons in Extended Data Figs. 6, 7). **c,** Anterior view of reconstructed TMEcen-neuron (tangential <u>cen</u>trifugal neuron of the <u>me</u>dulla) with ramifications in medulla and central brain. **d, e,** Cell and model responses for two TMEcen-neurons to flashed dark (**d**) and bright (**e**) bars. Two further TMEcen-neurons in Extended Data Figs. 8d-f. Response field header with outcome of two-way-ANOVA with “L” (“R”) being significant left (right) eye input and “I” significant interaction term (see Extended Data Table 1), otherwise “ns” meaning not significant.

If COcom only relayed information from one eye to the other they would be monocular. However, 5 of the 6 COcom-neurons were clearly binocular (Fig. 2d,f,g,i,j; Extended Data Table 1). A possible neuronal circuit comprises a COcom-neuron transmitting visual information to the contralateral optic lobe, but also receiving information from the contralateral eye via other COcom-neurons, and in this way generating its binocularity (see below).

Four of the binocular COcom-neurons showed evident binocular interactions in the central parts of the response fields (at 15-100mm distance and ±20° eccentricity; Fig. 2d,f,g,j). Behavioural experiments^6^ have shown that mantis stereopsis operates for prey capture over this region of 3D-space. Three neurons had well-localised excitatory peaks (Fig. 2d,f,g) for a preferred 3D location. These were well modelled by combining binocular excitation at the preferred location with inhibition in peripheral regions (Extended Data Fig. 4 a,b,c). In the fourth neuron the central region was void of excitation, because of inhibition by input from the contralateral eye in the centre of the visual field (Fig. 2 j,k). In vertebrates such cells are known as “tuned-inhibitory neurons”, in contrast to “tuned-excitatory neurons” whose receptive fields have a similar structure in both eyes^22^. The neuron from Fig. 2a,b,c,d also showed disparity tuning to the spiralling disc stimulus (Wilcoxon rank-sum test p=0.0043, Fig. 2e).

**Figure 4.**
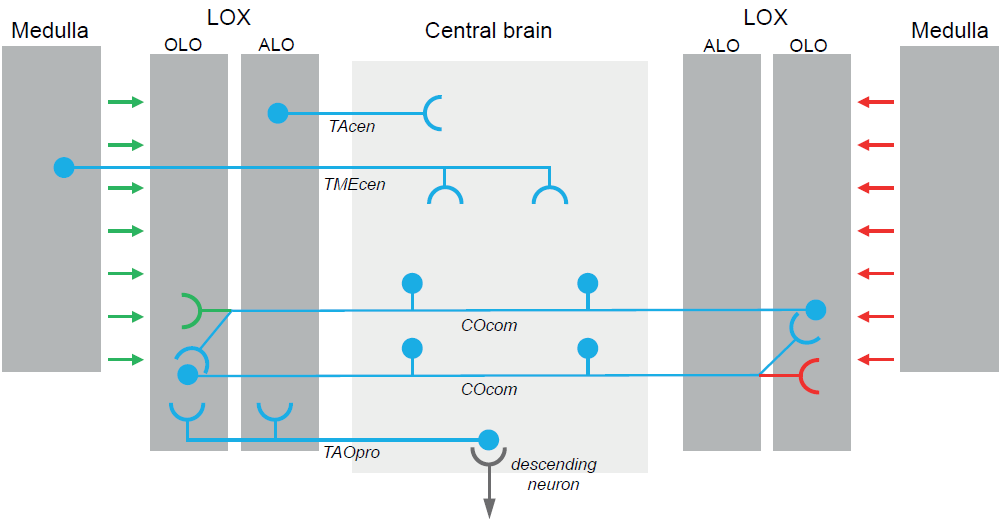
Neuronal circuitry for binocular disparity in the praying mantis brain. Grey shaded areas represent brain regions with ramifications of the recorded neurons; optic lobe regions are shaded darker. Feedforward visual information flow from left eye is illustrated in green, from right eye in red and combined, binocular blue. Hypothetical descending neuron is shown in grey. Input regions are shown with semi-circles, output with disks. For clarity TMEcen and TAcen are shown only on left side.

The morphology of two additional, disparity-sensitive neuron types suggests that they convey information centrifugally (soma in central brain - Fig. 3a,c; beaded terminal neurites in optic lobe - Extended Data Fig. 5) from the central brain to the LOX (*TAcen-neurons*) and the medulla (*TMEcen-neurons*). All 5 TAcen-neurons tested with bright bars were clearly disparity tuned (Fig. 3b, Extended Data Figs. 6). They have broad excitatory receptive fields with peak response for far distances or even diverging lines of sight. Thus, TAcen-neurons are suited to provide information about the distant, bright visual background in front of which dark prey-targets are detected by other classes of neuron like COcom.

All 4 TMEcen-neurons were also disparity tuned (Fig. 3d,e; Extended Data Fig. 8). However, unlike TAcen-neurons, different TMEcen-neurons responded to different disparities and even contrasts, comprising preferences for dark objects in catching range (Fig. 3d) or far away (Extended Data Fig. 8e) and bright objects at diverse distances (Fig 3e and Extended Data Fig. 8b,d,f). The example neuron response shown in Fig. 3e has a diagonal excitatory structure, meaning the cell responded to object distances independent of the azimuthal direction leftwards from the midline out to as far as we measured (32°). Less extensive diagonal structure was also seen in two COcom neurons (Fig 2d,g). This diagonal structure is a hallmark of vertebrate *complex cell* responses, which are believed to result from converging simple cell units^10^.

Figure 4 summarises the 4 neuronal classes of disparity-tuned insect neurons presented here. A working hypothesis consistent with our data is that disparity sensitivity for dark prey-like objects is established in the LOX by brain-spanning COcom neurons which transmit information between the outer lobes in each hemisphere. Disparity information is then passed on to projection neurons like TAOpro and hence to descending neurons which deliver the signal for the predatory strike.

TAOpro also receives input in the anterior lobe which is the output region for centrifugal TAcen neurons, tuned to bright background in front of which dark prey moves. TAcen neurons thus could modulate the gain of TAOpro-neurons, potentially preventing strikes if too much background is occluded (e.g. by a large object like a bird). Finally, centrifugal TMEcen neurons transmit disparity information from the central brain to the medulla, the early (second) visual neuropil. Here TMEcen could either boost overall neuronal processing when relevant stimuli occur, or it could guide attention in 3D-space. Insect centrifugal neurons have been repeatedly shown to modulate visual processing including involvement in selective attention^23–27^.

Our work rules out speculations that disparity sensitive neurons are not present in the praying mantis brain at all^9^ or occur only in a very late step, fusing pre-processed monocular signals in the central brain^6,9^. In a compelling vindication of the “retinal disparity hypothesis”, we have identified neurons which are tuned to different locations in 3D-space, at a range of distances and horizontal eccentricities, confirming disputed deductions from behavioural data^4,6^. A model proposed to explain disparity selectivity in the vertebrate visual cortex^10^ also captures the behaviour of most mantis neurons. Binocular disparities are computed early in the visual pathway, in the lobula complex, and fed back even earlier, to the medulla. Disparity feedback has not been reported in vertebrates nor included in machine stereo algorithms, but its discovery in insect vision implies it plays an important role. Thus, insect stereopsis suggests new approaches to machine stereo vision and demands new reflection regarding stereo vision in humans.

## Methods

### Animals

Experiments were carried out on adult praying mantids of the species *Hierodula membranacea* and *Rhombodera megaera*. Animals were housed in individual containers at a temperature of 25°C and a 12 h light/dark cycle. Adult animals were fed with a live cricket twice and younger mantids 3 times a week.

### Animal preparation

Animals were mounted on custom-made holders with BluTack® and wax; their head and mouthparts were immobilized by wax. A hole was cut into the posterior head capsule to allow access to the brain. Fat and muscle tissue surrounding the brain were removed. The neural sheath was stripped away at the region where the recording electrode was inserted. The gut was removed within the head capsule and prevented from leaking within the thorax by ligating it. A wire platform supported the brain from anterior to further stabilize it. During recording of neural activity the brain was submerged in cockroach saline.

### Neuronal recordings

We recorded intracellularly from 13 neurons with ramifications in the lobula complex and 4 with ramifications in the medulla. The neurons were identified by stainings with neuronal tracer (see below). Each cell was recorded in an individual animal. Microelectrodes were drawn from borosilicate capillaries (1.5 mm outer diameter, Hilgenberg, Malsfeld, Germany) on a microelectrode puller (P-97, Sutter Instrument, Novato, CA). Electrode tips were filled with 4% Neurobiotin (Vector Laboratories, UK) in 1 M KCl and their shanks with 1 M KCl. The electrodes had tip resistances of 70–150 MOhm. Signals were amplified (BA-03X amplifier; NPI), digitized (CED1401 micro; Cambridge Electronic Design, UK), and stored using a PC with Spike2 software (Cambridge Electronic Design, UK). About 0.1-1 nA of depolarizing current was applied for several minutes to iontophoretically inject Neurobiotin immediately after recording and in some recordings in-between the stimulus sequences. We only injected and analysed those neurons for which we could acquire responses to the presentation of at least 10 repetitions of the bar stimulus.

### Histology

After neuronal recordings animal heads were fixed overnight in a mixture of 4% paraformaldehyde, 0.25% glutaraldehyde, and 0.2% saturated picric acid in 0.1 M phosphate buffer. Afterwards brains were dissected out of the head capsule. The labelled neurons were made visible for confocal laser scanning microscopy (Leica TCS-SP5 / SP8; Leica Microsystems) by treatment of the brains with Cy3-conjugated streptavidin (Dianova, Hamburg, Germany) as previously described^13^.

### Visual stimulation

We used anaglyph technology^3,12^ to present 3D stimuli on a computer monitor (DELL U2413 LED). Tethered mantids watched the computer screen through spectral filters while we performed neuronal recordings in their brain. We presented stimuli with different colours (green and blue) that matched the spectral properties of the filters so that each eye saw only the image it was intended to see. The computer screen was positioned at a viewing distance of 10 cm from the praying mantis.

All stimuli were custom written in Matlab (Mathworks) using the Psychophysics Toolbox^28–30^. We presented two main stimuli for the current study. Most importantly we analysed monocular and binocular response fields of neurons with a flashed bar stimulus. For this we divided the region of binocular overlap into 6 non-overlapping vertical stripes of 12.8° horizontal and 99.5° vertical extent (Fig. 1f). In this way we covered almost 77° of the fronto-azimuthal visual field. This is slightly wider than the approximately 70° binocular overlap of praying mantids^6^. Bars were presented either to one eye only, for recording monocular response fields, or two bars concurrently, one for the left and one for the right eye, for determining binocular response fields.

We used bars instead of structures with smaller vertical extent because of the comparatively short recording times possible with sharp electrodes. In this way we avoided the need to identify receptive field elevation while enabling us to vary horizontal disparity, the difference in the bar’s location between left and right eyes. Because insect eyes are offset horizontally and fixed on the head, horizontal disparity along with visual direction specifies a unique 3D position in space^5,31^, as shown in Fig. 1f. All bar combinations, including both monocular and binocular conditions, were shown in pseudorandom order. The bars were displayed for 250 or 500 ms with a pause of the same duration in between each presentation. After all bar positions had been displayed a pause of 1.7-4.5 s followed, before the procedure started again.

The second stimulus was similar to what was found earlier to be a very effective elicitor of the praying mantis prey capture strike^3^. A 22°-diameter dark disc in front of a bright background appeared peripherally and spiralled in towards the centre of the screen (Fig. 1d). On reaching the screen centre, after 5 s, it stayed there for 2 s before vanishing. Small quivering movements were superimposed on the principal spiral trajectory and in the final 2s stationary disc phase. The disc was simulated to float at 25 mm distance in front of the praying mantis in order to simulate an attractive target in catch range of the animal. This was achieved by presenting one disc on the left hand side, which was only visible to the right eye and a disc of identical dimensions slightly shifted to the right, which was only visible to the left eye. We refer to this stimulus condition as the “near” condition. As a control condition, the left and right eye discs were swapped so that the right eye now saw the right hand side disc and the left eye saw the left hand side disc (cf Extended Data Fig. 1c vs d).

### Microscopy and image data analysis

Whole mounts were scanned with confocal laser scanning microscopes (CLSM, TCS SP5 and SP8, Leica Microsystems, Wetzlar, Germany) with a 10x oil immersion objective lens (SP5) or a 10x or 20x dry lens (SP8). The detail scans of the neuritic endings in Fig. 2b and c were done with a 63er glycerol immersion objective lens and the SP5 microscope. The SP5 microscope was located in the Biology Department of Marburg University (Germany) and the SP8 microscope in the Bioimaging Unit at Newcastle University (UK).

Neuronal reconstructions were done with the SkeletonTree tool within Amira^32^. The reconstructed neurons were registered manually into a reference lobula complex^13^ in Amira 5.33. The schemes of the mantis brain were done in Adobe Illustrator CS5 (Adobe Systems, Ireland).

### Data evaluation

Data analysis was done in Matlab (The MathWorks, Natick, MA). Bar stimulus induced spike counts were determined in 250 ms time windows starting at time 1 ms when a bar was displayed. The background spike count was determined in 800 ms time windows preceding each stimulus sequence. Responses were converted to spiking rates per second and normalized by dividing by the highest spiking rate that occurred during either bar presentation or background firing, depending which one was higher. Afterwards the responses were averaged across identical stimulus conditions for each cell.

For several TAcen-neurons we also determined dark bar off-responses in a 200 ms time window starting 50 ms after the respective bar was switched off (Extended Data Fig. 7 and Extended Data Table 1).

We interpolated all binocular response fields from 6×6 to 100×100 with the Matlab function “imresize” in bicubic mode. An example raw plot and its upsampled version is shown in Extended Data Fig. 2.

Neuron rr170403 was only weakly stained and it was not possible to trace the main neurite into the central brain. Moreover, in the confocal scan it was partly superimposed by a second even weaker stained projection neuron. We included rr170417 in our analysis, because we consider it most likely to belong to the TAcen-class of neurons as identified by its typical ramifications in the anterior lobe of the lobula complex.

### Statistical analysis

Responsiveness of neurons to left or right eye stimulation was determined by 2-way-ANOVA (anova2-function in Matlab; requirement for significance p<0.05). The two factors were the location of the bar in the left and right eye respectively. Each factor had 7 levels, corresponding to the 6 possible bar locations plus the blank-screen condition. A significant main effect of each factor therefore means that the response differed between at least two different bar positions for the respective eye, and/or the response differed for at least one bar location from the spontaneous rate. A non-significant interaction term means that binocular response was well described by the sum of monocular responses; a significant interaction means that they combine non-linearly.

Responsiveness to the spiralling disc stimulus was determined for a selection of neurons via two-sided Wilcoxon rank sum test (Matlab ranksum-function; p<0.05) by comparing spike counts within a time window of 3.5s, starting 1.5 s after stimulus onset, between the near and control conditions.

### Modelling

For simulating response fields we applied a linear-nonlinear model developed for vertebrate stereopsis^10^. The model assumes that visual stimulation contributes excitatory or inhibitory input dependent on the eye and location of the stimulation; that is, the model contains receptive fields for both the left and the right eye (Fig. 1h). The inputs from both eyes are filtered by the receptive field and then summed linearly along with a tonic input, necessary to account for a non-zero background rate in some neurons. If the result is negative or zero, the mean response is zero. If the result is positive, the mean response is given by its value raised to some exponent. The value of the exponent, the tonic input, and the 6 azimuthal sub-regions of the left and right eye receptive fields, together form 14 free model parameters which we fitted to the mean neuronal response in 49 conditions (no visual stimulation, 12 monocular conditions and 36 binocular). The estimated receptive fields with azimuthal sub-regions of 12.8° width and 99.5° vertical extent obviously have coarse spatial resolution, however, they nevertheless provided valuable information about responsiveness of the neurons below spiking threshold. This information could not be gained from neuronal recordings directly, since the recordings usually did not reveal much information about subthreshold responses, presumably because recording sites were spatially distant from synaptic input sites. The only binocular neuron with major graded membrane potential changes was the TAOpro-neuron. However, analysis of subthreshold activity did not reveal the information our model receptive fields provided us with about inhibition provided by the right eye in the periphery of the frontal visual field. This might indicate that these inhibitory processes are not taking place within TAOpro itself but happen at an earlier stage. Our model receptive fields also capture processes that occur presynaptic/upstream to the recorded neurons.

## Acknowledgements

This work was funded by a Research Leadership Award RL-2012-019 from the Leverhulme Trust to JR. We thank Dr Sasha Gartside and Dr Richard McQuade for providing equipment and advice when setting up the laboratory, Prof Uwe Homberg, Dr Claire Rind, Dr Peter Simmons, Prof Yoshifumi Yamawaki, Dr Sid Henriksen, Prof Keram Pfeiffer, Prof Ignacio Serrano-Pedraza and Prof David O’Carroll for helpful discussions, Prof Geraldine Wright for lending a micromanipulator, Dr Alex Laude and Dr Rolando Palmini for help with the confocal microscopy, Prof Ignacio Serrano-Pedraza for providing spectral filters, and Dr Bruce Cumming, Dr Vivek Nityananda, and Prof Geraldine Wright for helpful comments on the manuscript.

## Author contributions

RR and JR conceived the aim of the study and the visual stimuli. RR built the electrophysiology laboratory, did all experiments, analysed the data and prepared the figures. JR programmed the model. RR and JR wrote the paper. JvH reconstructed the neurons and neuropils and prepared anatomical figure panels. GT programmed the stimuli and helped with synchronisation of visual stimuli and neuronal recordings.

---

**Extended Data Fig. 1.**
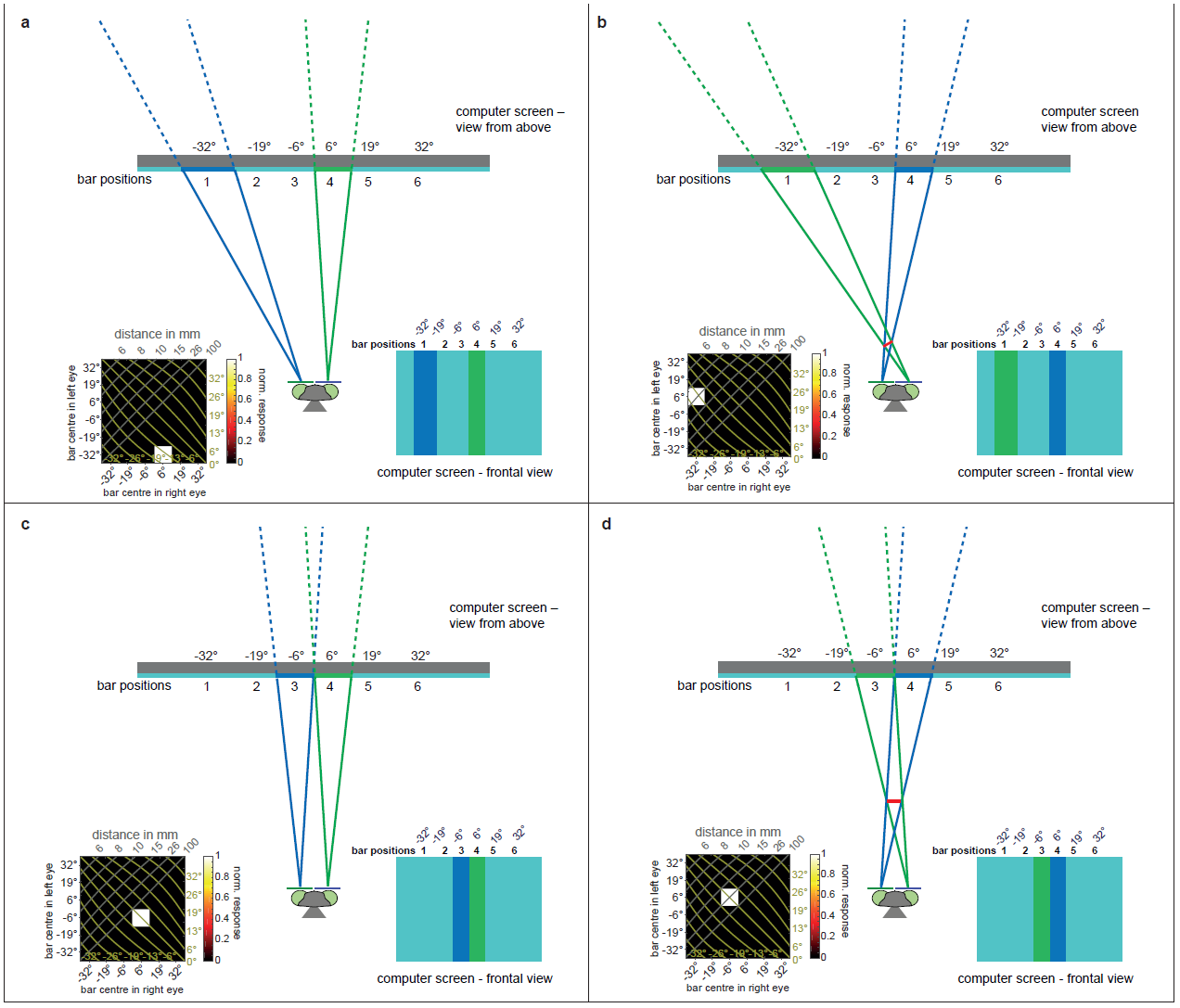
Visual stimulation and response field plotting in detail. **a-d**, Praying mantis with anaglyph filters watches computer screen with bar stimulus. Bottom left - binocular response field plot with neuronal response as would be observed if neuron responded exclusively to the shown stimulus configuration. Figures are not to scale. **a,c,** Lines of sight for left and right eye bar don’t intersect in front of screen (“uncrossed disparity”, control condition). These stimulus configurations are represented by points in the lower-right half of the binocular response plots. **b,d,** Lines of sight for left and right eye bar do intersect in front of screen (“crossed disparity”, near condition). These stimulus configurations correspond to a bar located in front of the screen (shown in red) and are represented by points in the upper-left half of the binocular response plots.

**Extended Data Fig. 2.**
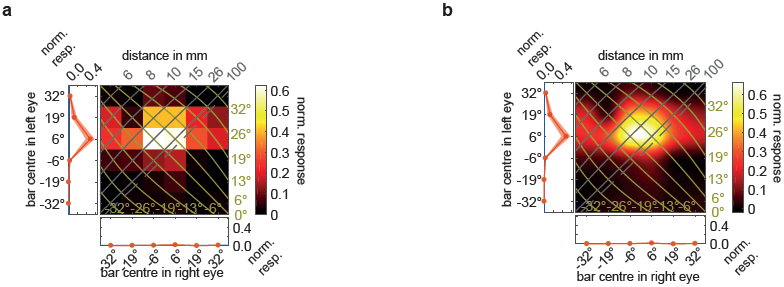
Raw and interpolated response field plots for TAOpro-neuron. **a,** Raw (non-interpolated) monocular and binocular response field plots. **b,** Interpolated binocular response field plot and non-interpolated, monocular response field plots.

**Extended Data Fig. 3.**
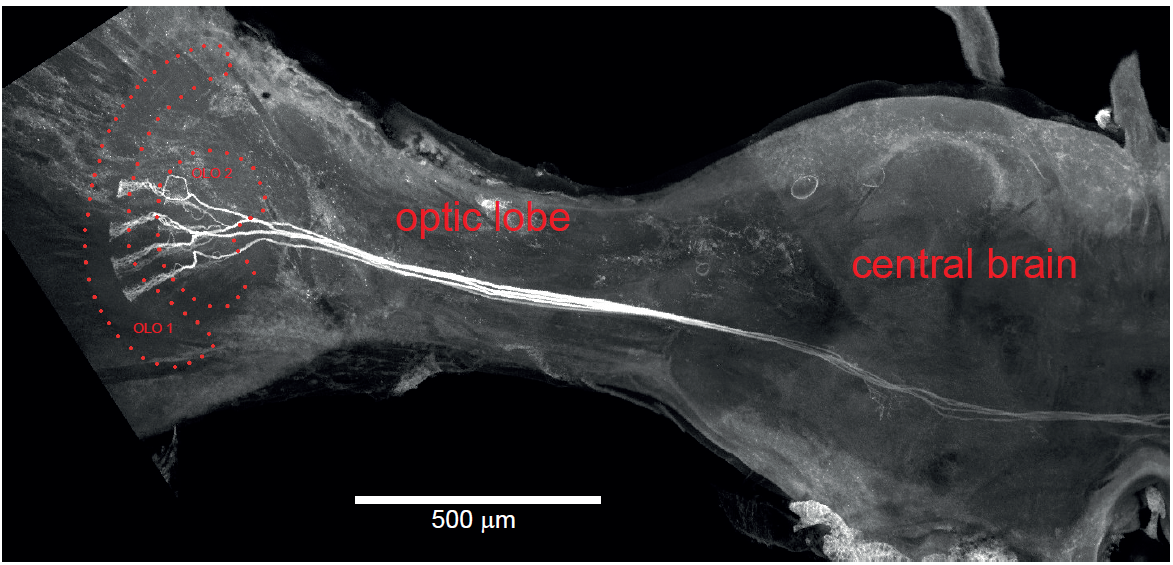
Array of COcom-neurons in the praying mantis brain. Maximum projection of confocal microscopy slices through left side of praying mantis brain (posterior view). Several COcom-neurons were stained during the experiment that culminated in the recording of neuron ID rr170117. OLO1, outer lobe 1 of lobula complex; OLO2, outer lobe 2 of lobula complex.

**Extended Data Fig. 4.**
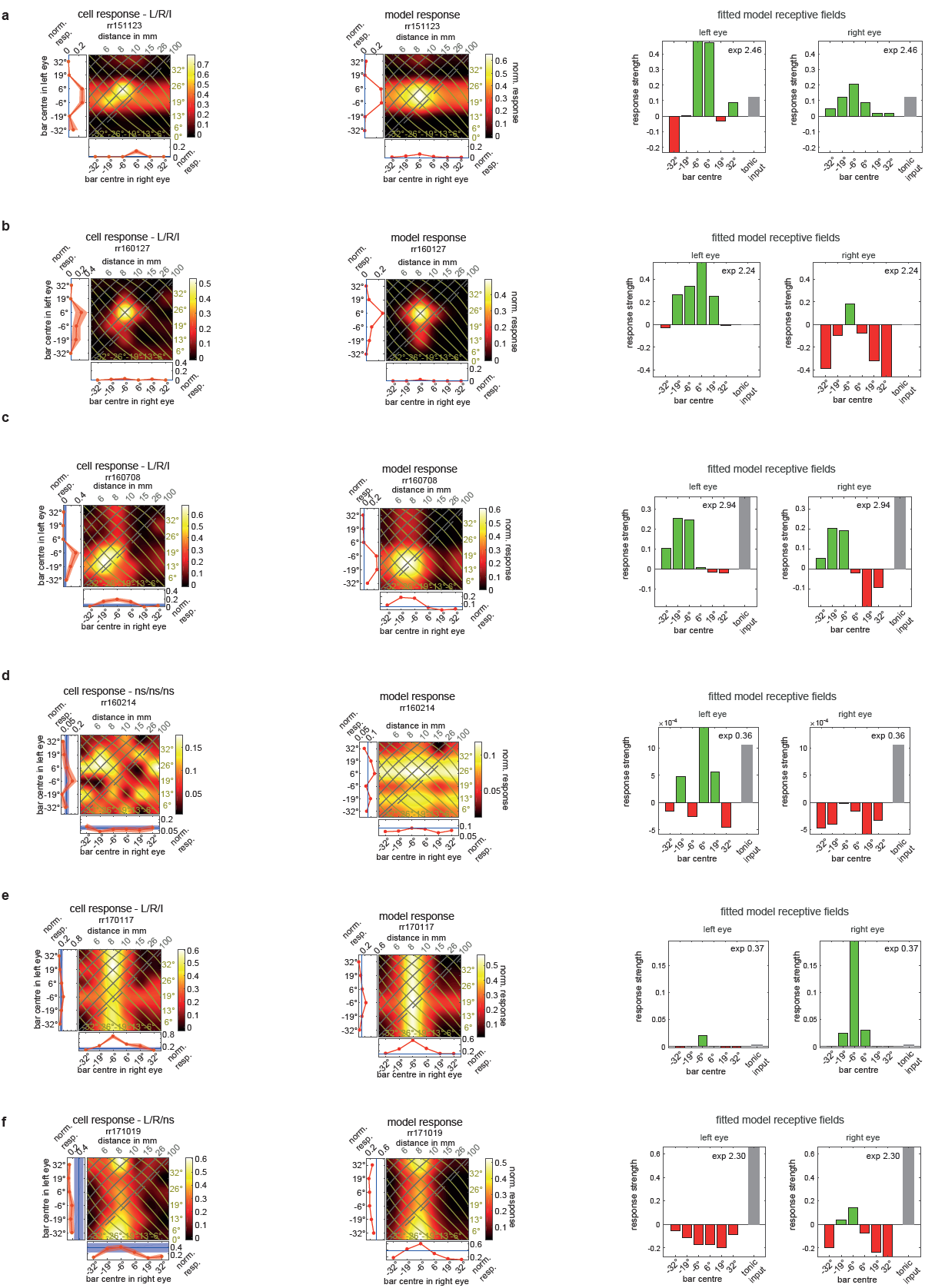
Response field plots and fitted receptive fields for COcom-neurons. **a-f,** Left panels show monocular and binocular response field plots for all recorded COcom-neurons as presented in Fig. 2. Response field headers state neuron ID and outcome of two-way-ANOVA with “L” (“R”) being significant left (right) eye input and “I” significant interaction term (see Extended Data Table 1), otherwise “ns” meaning not significant. Middle panels show response field plots for model predictions. Right panels show fitted receptive fields for left and right eye with excitations (green bars) and inhibitions (red bars) at corresponding azimuthal locations (x-axis). Grey bars show tonic input (shown in both RF plots but applied only once). Exponent in upper right corner (exp). Negative (positive) values on x-axis indicate locations left (right) of centre.

**Extended Data Fig. 5.**
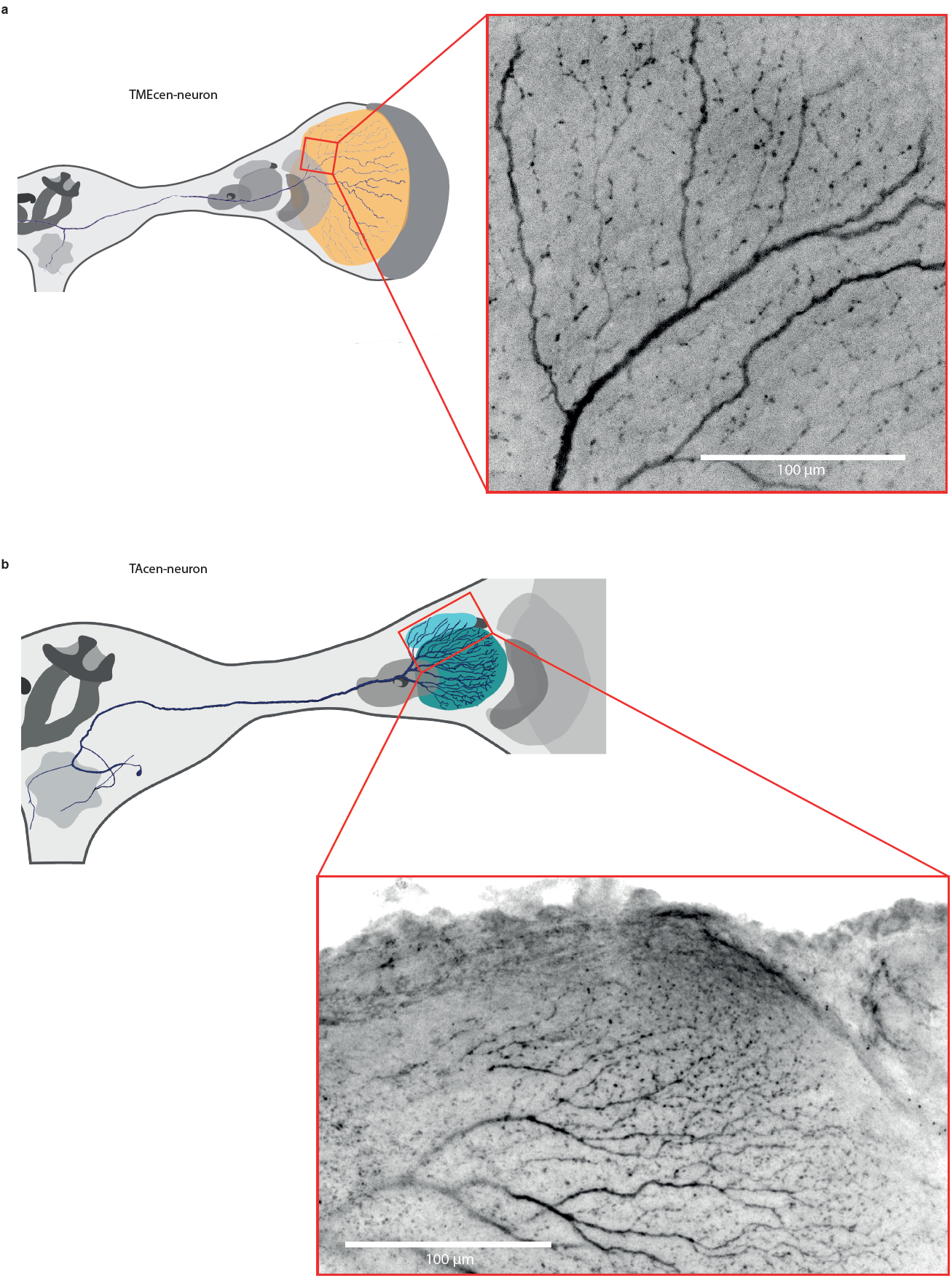
TAcen-and TMEcen-neurons have output arborizations in the optic lobe. **a,b,** Projection views of multiple confocal images for TMEcen-neuron (**a,** right) and TAcen-neuron (**b,** bottom right) branchings in left optic lobe. Location of recording windows indicated in left hand side schemes. In both neurons the terminal neurites are beaded/globular, indicating presynaptic regions^21^.

**Extended Data Fig. 6.**
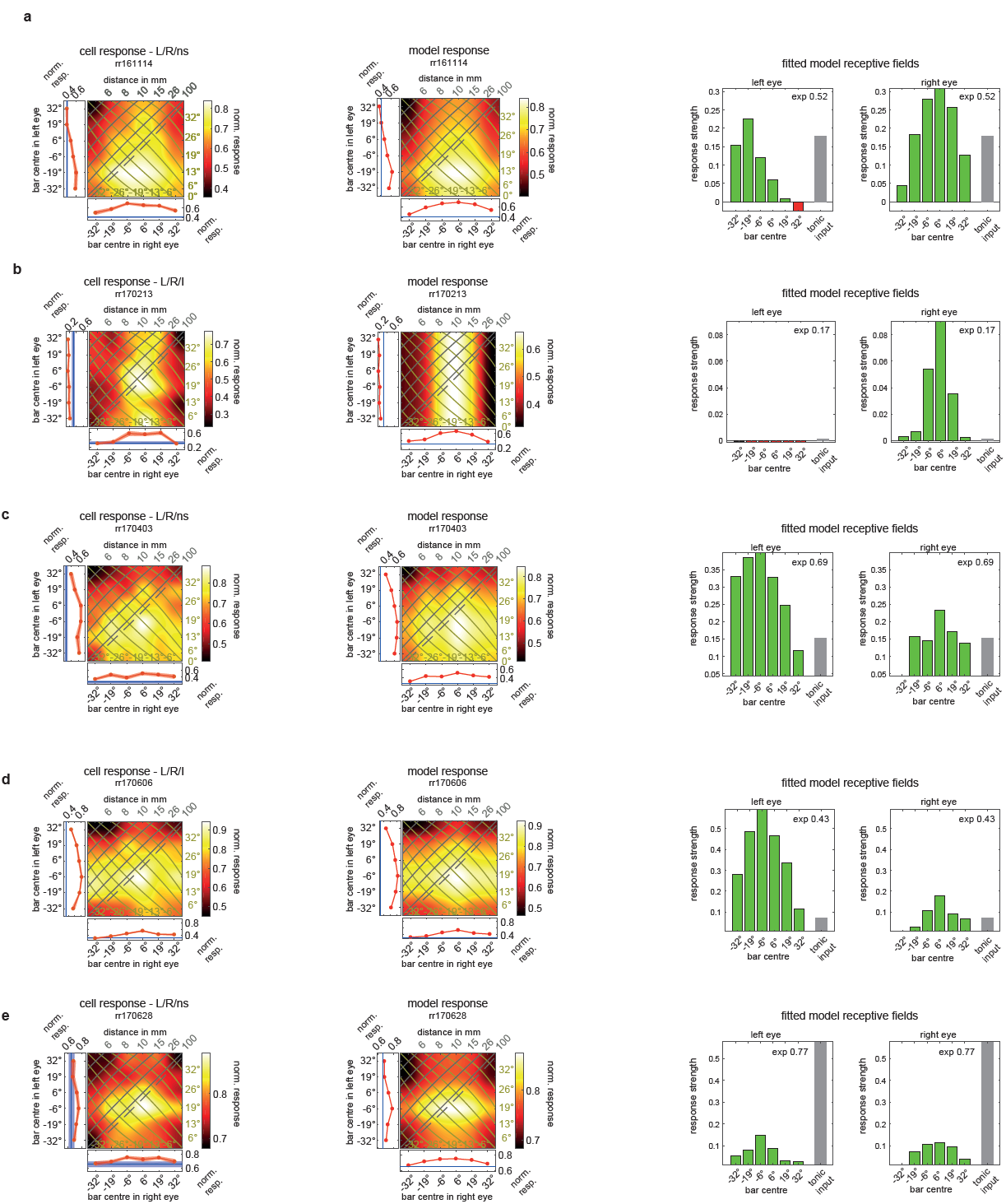
TAcen-neuron responses to flashed bright bars and model responses. **a-e,** Left panels show response field plots for TAcen-neuron bright bar stimulation. Response field headers state neuron ID and outcome of two-way-ANOVA with “L” (“R”) being significant left (right) eye input and “I” significant interaction term (see Extended Data Table 1), otherwise “ns” meaning not significant. Middle panels show raw response field plots for model predictions. Right panels show fitted receptive fields for left and right eye with excitations (green bars) and inhibitions (red bars) at corresponding azimuthal locations (x-axis). Grey bars show tonic input (shown in both RF plots but applied only once). Exponent in upper right corner (exp). Negative (positive) values on x-axis indicate locations left (right) of centre.

**Extended Data Fig. 7.**
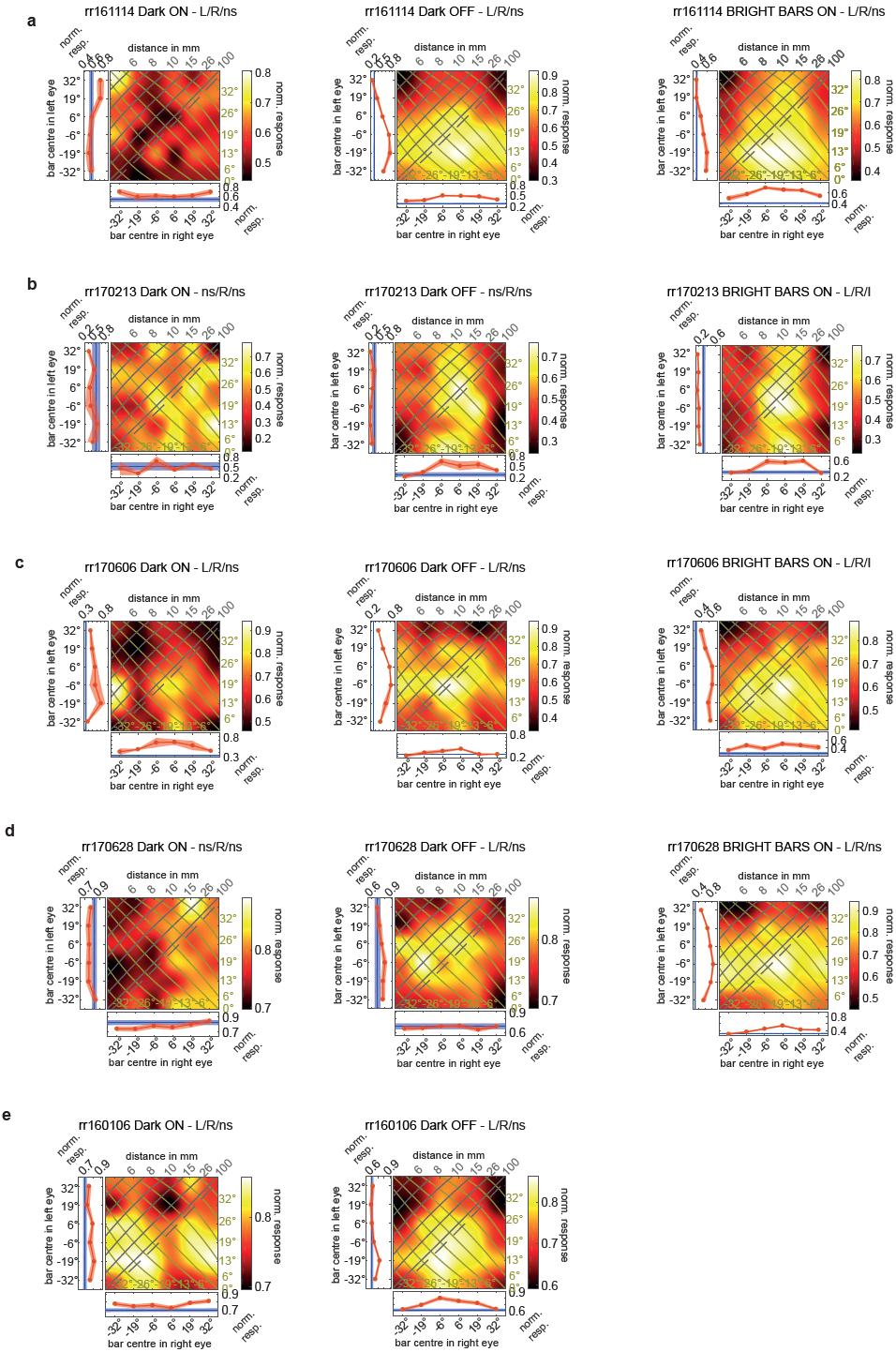
Dark bar on-and off-responses compared to bright bar on-responses in TAcen-neurons. **a-e,** Left panels show monocular and binocular response field plots for TAcen-neurons that have been tested for dark bar responses. Response field headers state neuron ID, stimulus type and outcome of two-way-ANOVA with “L” (“R”) being significant left (right) eye input and “I” significant interaction term (see Extended Data Table 1), otherwise “ns” meaning not significant. Left panels show dark bar on-responses (normalized spike count in 250ms time window starting at 0ms after stim onset), middle panels show dark bar off-responses (spike count in 200ms time window starting 50ms after dark bar vanished). **a-d,** Right panels show bright bar on-responses (normalized spike count in 250ms time window starting at 0ms after stim onset).

**Extended Data Fig. 8.**
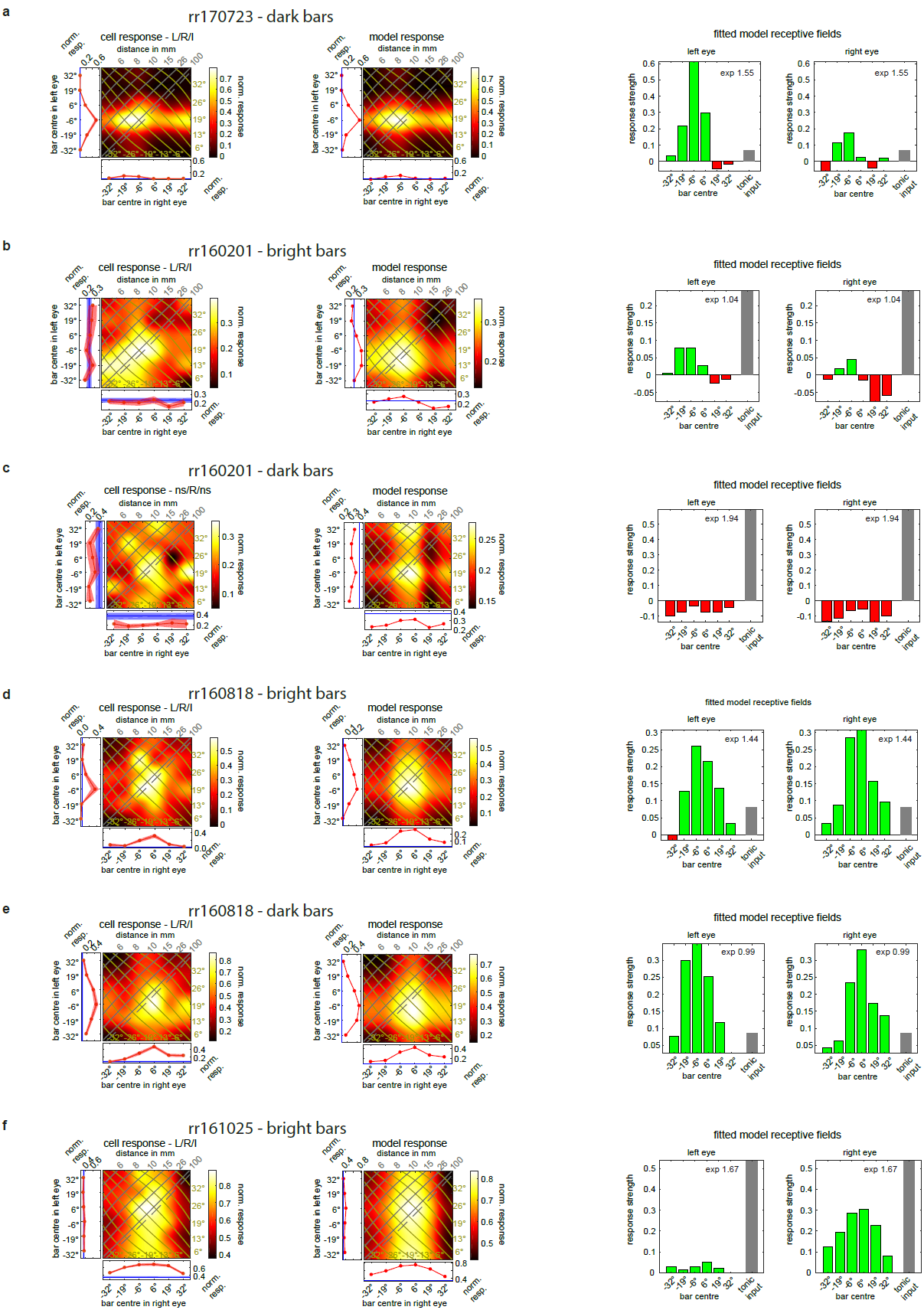
Response field plots and fitted receptive fields for TMEcen-neurons. **a-f**, Left panels show monocular and binocular response field plots for four TMEcen-neurons. Response field headers state neuron ID and outcome of two-way-ANOVA with “L” (“R”) being significant left (right) eye input and “I” significant interaction term (see Extended Data Table 1), otherwise “ns” meaning not significant. Middle panels show model predictions and right panels fitted receptive fields for left and right eye with excitations (green bars) and inhibitions (red bars) at corresponding azimuthal locations (x-axis). Grey bars show tonic input (shown in both RF plots but applied only once). Exponent in upper right corner (exp). Negative (positive) values on x-axis indicate locations left (right) of centre.

**Extended Data Table. 1.** | Responsiveness of neurons to left and right eye bar stimulation. 6th column indicates whether there was a significant (p<0.05) main effect of left (L) or/and right (R) eye stimulation on neuronal response as tested with 2-way-ANOVA (see Methods: Statistical analysis). A significant interaction term in the 2-way-ANOVA is indicated by “I”, non-significance by ‘ns’. Degrees of freedom were 6 for left and right eye main effects, respectively and 36 for interaction terms. F – and p-values are provided in 7^th^ and 8^th^ column, respectively. p-values smaller than 0.0001 are given as 0. Recordings were usually done without concurrent depolarizing current injection unless indicated by ‘inj’ with the amount of current indicated in nA. When trials with and without current injection were included it is stated with ‘w/o inj’. On-responses were evaluated in a 250 ms time window starting 1 ms after stimulus onset. Off responses were evaluated in a 200 ms time window starting 50 ms after stimulus off. *Hierodula, Hierodula membranacea*; ND, not determinable (no clear peak response); OL, optic lobe; CB, central brain; *Rhombod., Rhombodera megaera*.

| Neuron class | Neuron ID | Input > output | Stimulus type | Repetitions | ANOVA | F-values | p-values | Species |
| --- | --- | --- | --- | --- | --- | --- | --- | --- |
| TAOpro | rr160427 | Left OL > CB | Dark bars (on) | 18 | L,R,I | 100.79 / 16.67 / 3.73 | 0 / 0 / 0 | *Hierodula* |
|  | rr160427 |  | Bright bars (on) | 3 | L,ns,ns | 13.83 / 1.76 / 0.89 | 0 / 0.1154 / 0.6505 |  |
| COcom | rr151123 | Left OL > right OL | Dark bars (on) | 11 | L,R,I | 93.04 / 5.33 / 2.93 | 0 / 0 / 0 | *Hierodula* |
| COcom | rr160127 | Right OL > left OL | Dark bars (on) | 10 | L,R,I | 7.76 / 6.24 / 2.14 | 0 / 0 / 0.0002 | *Hierodula* |
| COcom | rr160708 | ND | Dark bars (on) | 11 | L,R,I | 21.38 / 23.47 / 2.43 | 0 / 0 / 0 | *Hierodula* |
| COcom | rr160214 | Right OL > left OL | Dark bars (on) | 29 | ns,ns,ns | 1.88 / 0.36 / 1.35 | 0.0813 / 0.9053 / 0.0814 | *Hierodula* |
|  | rr160214 |  | Bright bars (on) | 12 | ns,R,ns | 0.13 / 2.95 / 0.72 | 0.993 / 0.0076 / 0.888 |  |
| COcom | rr170117 | Right OL > left OL | Dark bars (on) | 10 | L,R,I | 4.51 / 76.63 / 2.02 | 0.0002 / 0 / 0.0006 | *Rhombod.* |
| COcom | rr171019 | Right OL > left OL | Dark bars (on) | 13 | L,R,ns | 2.62 / 26.02 / 1.15 | 0.0162 / 0 / 0.2509 | *Hierodula* |
|  | rr171019 |  | Bright bars (on) | 4 (0.2nA inj) | ns,ns,ns | 1.45 / 1.43 / 1.35 | 0.1978 / 0.2081 / 0.1079 |  |
| TAcen | rr161114 | CB > left OL | Bright bars (on) | 15 | L,R,ns | 39.08 / 69.01 / 0.65 | 0 / 0 / 0.9456 | *Rhombod.* |
|  | rr161114 |  | Dark bars (on) | 12 | L,R,ns | 4.67 / 3.12 / 1.43 | 0.0001 / 0.0052 / 0.0524 |  |
|  | rr161114 |  | Dark bars (off) | 12 | L,R,ns | 162.31 / 40.94 / 1.26 | 0 / 0 / 0.1501 |  |
| TAcen | rr170213 | CB > left OL | Bright bars (on) | 12 | L,R,I | 3.83 / 94 / 1.68 | 0.001 / 0 / 0.009 | *Rhombod.* |
|  | rr170213 |  | Dark bars (on) | 4 (0.4nA inj) | ns,R,ns | 0.89 / 2.68 / 1.1 | 0.5039 / 0.0171 / 0.3374 |  |
|  | rr170213 |  | Dark bars (off) | 4 (0.4nA inj) | ns,R,ns | 1.39 / 23.53 / 0.76 | 0.2229 / 0 / 0.8359 |  |
| TAcen | rr170606 | CB > left OL | Bright bars (on) | 10 (w/o 0.1nA inj) | L,R,I | 249.17 / 21.76 / 2.16 | 0 / 0 / 0.0002 | *Hierodula* |
|  | rr170606 |  | Dark bars (on) | 2 (0.1nA inj) | L,R,ns | 8.56 / 7.39 / 0.91 | 0 / 0 / 0.6135 |  |
|  | rr170606 |  | Dark bars (off) | 2 (0.1nA inj) | L,R,ns | 42.85 / 5.8 / 0.59 | 0 / 0.0001 / 0.9486 |  |
| TAcen | rr170628 | CB > left OL | Bright bars (on) | 17 | L,R,ns | 26.21 / 23.21 / 0.66 | 0 / 0 / 0.9357 | *Rhombod.* |
|  | rr170628 |  | Dark bars (on) | 13 (w/o 0.3nA inj) | ns,R,ns | 0.97 / 6.51 / 0.62 | 0.445 / 0 / 0.9594 |  |
|  | rr170628 |  | Dark bars (off) | 13 (w/o 0.3nA inj) | L,R,ns | 17.3 / 3.41 / 0.76 | 0 / 0.0026 / 0.8385 |  |
| TAcen | rr170403 | CB > left OL | Bright bars (on) | 11 | L,R,ns | 61.34 / 16.6 / 0.92 | 0 / 0 / 0.6042 | *Hierodula* |
| TAcen | rr160106 | CB > left OL | Dark bars (off) | 13 | L,R,ns | 9.3 / 4.59 / 0.69 | 0 / 0.0001 / 0.9122 | *Hierodula* |
|  | rr160106 |  | Dark bars (on) | 13 | L,R,ns | 23.35 / 39.57 / 0.79 | 0 / 0 / 0.8016 |  |
| TMEcen | rr160201 | CB > left OL | Bright bars (on) | 32 | L,R,I | 6.45 / 6.1 / 1.49 | 0 / 0 / 0.0325 | *Hierodula* |
|  | rr160201 |  | Dark bars (on) | 15 | ns,R,ns | 0.7 / 2.22 / 0.83 | 0.6468 / 0.0391 / 0.7516 |  |
| TMEcen | rr160818 | CB > left OL | Bright bars (on) | 13 | L,R,I | 23.18 / 30.26 / 2.1 | 0 / 0 / 0.0003 | *Hierodula* |
|  | rr160818 |  | Dark bars (on) | 15 | L,R,I | 86.25 / 60.58 / 1.68 | 0 / 0 / 0.0082 |  |
| TMEcen | rr161025 | CB > left OL | Bright bars (on) | 15 | L,R,I | 6.49 / 189.94 / 1.54 | 0 / 0 / 0.0247 | *Hierodula* |
| TMEcen | rr170723 | CB > left OL | Dark bars (on) | 23 (0.15nA inj) | L,R,I | 288.74 / 25.96 / 2.95 | 0 / 0 / 0 | *Rhombod.* |

